# The second life of collection - how the scientific value of natural history holdings can be deliberately increased

**DOI:** 10.64898/2026.09.16.752043

**Authors:** Jerzy Błoszyk, Marcin Lawenda

## Abstract

Natural history collections cannot be assessed by the size of their holdings, because their scientific value is a multidimensional and dynamic property that can be deliberately shaped. The authors argue that this value can be systematically increased through purposeful curatorial and research actions, both without acquiring new material and through its continuous replenishment. We propose an analytical framework for describing the scientific value of a collection, based on five dimensions: epistemic distinctiveness, data integrity and accessibility, documented and potential research applications, unique taxonomic status, and compliance with ethical and legal standards. The framework is complemented by a dynamic perspective describing the conditions under which the value of a collection rises and falls over time. We then present a typology of actions that increase collection value incrementally, distinguishing four categories according to the object of the action: (i) enriching the data record, (ii) enriching the physical specimen, (iii) establishing formal status, and (iv) increasing visibility and interoperability. Each category of actions is linked to the corresponding dimensions of scientific value, so that the typology and the framework together form a single analytical tool. Both perspectives are illustrated with two collections deposited in the Natural History Collections of Adam Mickiewicz University in Poznań. The first is an acarological collection of soil samples together with its derivatives, comprising collections of nomenclatural types, microscope slides, SEM images and molecular sequences deposited in global gene banks. The second is a conchological and virtual collection of the Roman snail (Helix pomatia), which illustrates the transition from a set of physical specimens to a resource of spatial digital data. The proposed approach provides practical tools for planning collection development, for communicating the importance of collections to funding bodies, and for identifying the untapped research potential of existing holdings.

## 1 Introduction

Collecting natural history specimens has a long history. Assembled for a variety of motives, natural history holdings have undergone a distinctive metamorphosis, from cabinets of curiosities to modern museum institutions housing millions of specimens that document the diversity of life on Earth. Whatever their origin, museum collections are once again gaining importance in the eyes of naturalists, and in ways their creators could not have foreseen. What, then, are these holdings, and why are they becoming important in twenty-first-century science? What role can stuffed animals, entomological collections locked in display cases, invertebrates preserved in alcohol or plants dried in herbaria play in modern biology, dominated by molecular research and the analysis of large data sets? Answering these questions is the subject of this article.

A special place among the applications of natural history collections belongs to faunistics, the branch of biology concerned with the composition and distribution of fauna in a given area and at a given time. Faunistic research makes it possible to assess the conservation status of threatened species, to monitor environmental change and to preserve biological diversity, and it is therefore one of the fundamental tools of sustainable management of natural resources. Natural history collections are to faunistics what archives are to history, but their importance extends far beyond faunistics alone, into taxonomy, molecular biology, ecology and biodiversity conservation. It is precisely this capacity of a collection to answer questions that were not yet being asked when it was assembled that constitutes the essence of what we call in this article the second life of a collection.

We argue in this article that the scientific value of a natural history collection is neither a fixed property nor a simple function of its size, but a complex and dynamic profile that can be deliberately shaped. After a synthetic review of the state of the art (Section 2), Section 3 discusses five dimensions of the scientific value of a collection, which together form the analytical framework for the argument, along with an account of how that value changes over time. Section 4 presents a typology of curatorial and research actions that increase this value incrementally. Section 5 illustrates both perspectives with two collections held in the Natural History Collections of Adam Mickiewicz University in Poznań: (i) the collection of soil samples together with its derivatives, and (ii) a collection, originally conchological, of the largest Polish land snail, the Roman snail (Helix pomatia). This is the only mollusc harvested from natural sites in Poland for commercial purposes, which entails specific requirements arising from the Nature Conservation Act.

## 2 State of the art

The importance of natural history collections as research infrastructure has been the subject of a growing number of review papers over the last two decades. The common denominator of this literature is the thesis that natural history collections enter the twenty-first century as a resource of increasing rather than decreasing importance. This follows, paradoxically, from the development of analytical methods, which makes it possible to put questions to historical material that had not yet been formulated when that material was being assembled [1], [2].

Works documenting the multidimensional character of the scientific value of collections remain a fundamental point of reference. Suarez and Tsutsui [1] were among the first to demonstrate that natural history collections generate value on many planes simultaneously, from taxonomic and evolutionary research to ecology, biodiversity conservation and the medical sciences, and that this cannot be captured by any single quantitative indicator. Miller et al. [2] developed this thesis in an operational direction, formulating requirements for the standards of collecting and documenting new specimens in a twenty-first-century perspective. Meineke et al. [3] and Holmes et al. [4] showed empirically that historical collections supply time-series data that make it possible to track species responses to climatic change and habitat transformation, and that this value grows retroactively with every new method that allows questions about the past to be asked.

A separate and rapidly developing strand is the literature devoted to digitisation as a tool for increasing the value of existing holdings. The notion of digitisation has evolved from the simple scanning of labels towards a much broader concept. Hedrick et al. [5] distinguished two stages of this process: Digitization 1.0, which converts the metadata of a physical specimen into accessible digital data, and Digitization 2.0, which creates digital workflows operating solely on digitised data and returning value to specimens through new layers of annotation, automated identification methods and global collaborative networks. Popov et al. [6] documented the economic and scientific effects of digitising the collections of the Natural History Museum in London, showing a measurable increase in citations and in new research applications as a direct consequence of making the data available in global aggregators. Wieczorek et al. [7] showed that standardising metadata to the Darwin Core schema is a necessary condition for such aggregation, and Wilkinson et al. [8] defined the FAIR principles as a normative framework for research data management, now a standard requirement of publicly funded projects.

In parallel, a literature developed on enriching collection holdings with molecular data. Hebert et al. [9] introduced DNA barcoding as a method of biological identification based on the sequencing of standardised genetic markers, and Ratnasingham and Hebert [10] built the BOLD data infrastructure linking sequences to the records of collection specimens. Holmes et al. [4] showed that the retroactive recovery of DNA from historical museum specimens opens a new view of evolutionary and phylogeographic data on a time scale inaccessible to field research. In the acarological context, the integration of morphology and DNA barcoding was tested on soil-dwelling and nest-dwelling mites of the Western Palaearctic. Close to 80% of species were correctly identified by molecular means, but an exact one-to-one correspondence with a BIN in the BOLD system was achieved for only about 60%. High intraspecific divergence in roughly one third of the species further indicated the presence of cryptic taxa [11].

The question of the nomenclatural status of specimens and its relation to the scientific value of collections was analysed by Sluys [12], who demonstrated that type specimens are an irreproducible and irreplaceable element in the architecture of biological nomenclature, and that their proper documentation and registration in systems such as ZooBank [13] is a condition of the long-term stability of taxonomy. Lendemer et al. [14] proposed the Extended Specimen concept as an architecture integrating all derivative data objects around a single identified specimen, now regarded as the target standard for the DiSSCo and iDigBio infrastructures.

The most recent literature adds two important qualifications to the picture drawn by earlier work. First, Forbes et al. [15], analysing more than 150 million GBIF records, showed that the rate at which new specimens are collected has been falling structurally for several decades, which means that the value of collections as global research infrastructure is threatened not only by curatorial neglect but also by declining collecting activity. Second, a formal treatment of collecting priorities in decision-theoretic terms has been proposed, integrating information value, information need and information cost into a unified framework for resource optimisation [16]. Both works confirm the direction of the argument advanced here while pointing to the possibility of quantitative formalisation, a direction this article deliberately does not pursue, adopting instead a qualitative and typological approach oriented towards the needs of practitioners and collection curators. Earlier work by some of the present authors [17] also belongs to the quantitative strand: it proposed a method for determining the relativistic value of a natural history collection (the Relativistic Collection Value). A set of assessment criteria was defined there, weights were assigned to them, and they were combined into an algorithm implemented as an electronic form that computes an aggregate value for the collection. The immediate motivation was to align maintenance subsidies with the actual value of the holdings. The present article is a complementary layer to that approach. Instead of aggregating criteria into a single number, it decomposes scientific value into dimensions and indicates the actions that raise each of them. It therefore answers not the question of how much a collection is worth, but of what to do to make it worth more.

A feature shared by the standards discussed above is that their natural unit of reference remains the individual specimen: Darwin Core organises its metadata, Extended Specimen links derivative objects to it, and the FAIR principles define the conditions under which records so constructed circulate. In parallel, although far more slowly, standards for describing collections as wholes are developing, from Natural Collections Descriptions to the Latimer Core standard ratified by TDWG in 2024 [18]. It allows a collection to be characterised through its taxonomic, geographic and temporal scope, without descending to the level of individual records. Since the subject of this article is the value of a collection rather than the value of a specimen, it is this second level of description that provides the proper background for the analytical framework proposed below.

## 3 The scientific value of natural history collections

Natural history museums hold some of the most information-rich archives of life on Earth (specimens, tissues, field data and environmental samples), forming research infrastructure of a unique and irreproducible character [1], [2]. The scientific value of a natural history collection is not, however, a one-dimensional notion, and it cannot be reduced to a simple set of figures or indicators. It is a complex property arising from several interdependent dimensions which together determine the extent to which a given collection can stimulate scientific discovery.

The framework proposed below emerged from combining the review presented above with the curatorial practice of the authors and with the set of assessment criteria developed while constructing the relativistic value of a collection [17]. The starting point was the aspects of value that recur in review papers, supplemented by two that are weakly represented there but important in daily work with a collection: the nomenclatural status of specimens and the ethical and legal compliance of the holdings. The resulting set of five dimensions was then tested against the two collections described in Section 5, taking as the closure criterion that every feature regarded as important by the curators of those holdings can be assigned to one of the dimensions. The dimensions are not mutually exclusive: the FAIR principles and persistent identifiers belong simultaneously to data integrity and to ethical and legal compliance, as discussed below. The framework is descriptive and ordering rather than metrical: it allows discussion of collection value to be organised and neglected areas to be identified, but it supplies neither a scale nor a numerical indicator. This is not an argument against quantification as such. A composite index built from weighted criteria, of the kind proposed in the work cited above, aggregates value into a single comparable figure. The framework below asks the complementary question of which components that figure combines and how each of them can be raised. The level of description at which the framework operates corresponds to the one organised by the Latimer Core standard: the taxonomic scope of a collection falls under unique taxonomic status, its geographic and temporal scope under epistemic distinctiveness, the completeness and format of its metadata under data integrity and accessibility, and its legal status under ethical and legal compliance [18].

The foundation of the scientific value of a collection is its **epistemic distinctiveness**, defined by the breadth and depth of material that cannot easily be replicated [1], [2]. A collection acquires this feature when it contains long series of specimens from defined sites, taken by comparable methods over many years, rich material from diverse environments, or carefully selected tissues appropriately preserved with future analytical techniques in mind. Such collections make comparisons across time and space possible, and they support both replication studies and meta-analyses. The scientific value of a collection also grows when methods and metadata are standardised sufficiently to allow specimens to be compared both within the collection and against external data sets. Temporal continuity, a transparent sampling design and well-documented gaps matter as much as the presence of unique taxa. Epistemic distinctiveness also includes the synergistic effect of sharing a collection: when the same material is used in parallel by many researchers working on different groups of organisms, each successive analysis raises the information value of the samples for the others as well, something that cannot be achieved in private holdings or in material assembled for a single project [3], [4], [19].

The next key dimension is **data integrity and accessibility**. Labels, field notes, georeferencing and persistent identifiers locate specimens in specific places, dates and research methods, while well-developed databases transform physical (analogue) holdings into digital resources that researchers actually use [8], [6]. Databases that provide access both to raw archival data and to contemporary data significantly extend the reach of a collection and the number of studies it inspires. Every collection holding exceptional biological material but lacking adequate documentation and cataloguing is an irreversible loss to science.

The third dimension is **documented and potential research applications**. The use made of a collection so far, measured by the number of doctoral theses, monographs, peer-reviewed articles, data sets and loans, is the best evidence of its real importance to science. The most direct beneficiaries of this kind of resource are faunistic and taxonomic research, for which voucher specimens together with their metadata are the only possible comparative material, followed by ecology, zoogeography, molecular biology and biodiversity conservation [20], [21]. The value of a collection is not, however, exhausted by its use to date. Structural features such as temporal scope, unique habitats and microhabitats, or the geographic region sampled point to research problems that the collection will help to solve in the future. This is tied to the development of new research methods, from molecular analyses through stable isotopes to computed microtomography [3], [4]. Appropriate storage conditions and protection against pests safeguard this potential, guaranteeing the stability and interpretability of the specimens held over the long term.

The scientific value of a collection is also determined by its **unique taxonomic status**. Type specimens (nomenclatural types), specimens of extinct or critically endangered species, and those representing endemics of restricted geographic range give a collection an exceptional and irreplaceable character. They are the indispensable basis of zoological and botanical nomenclature and the reference point for any systematic revision. As important as holding type specimens is the growth trajectory of the collection. An actively developing collection, enriched with new material and data, has greater potential than a closed one, although the latter may also be invaluable for historical and comparative analyses [22], [23].

A final and inseparable element of scientific value is **compliance with ethical and legal standards**. Transparent collecting permits, a policy on destructive sampling and rules for depositing material in accredited repositories determine the legality and sustainable use of the holdings. Without these elements, even the richest collection may be unavailable for international collaboration or may forfeit the trust of the scientific community. Compliance with the FAIR principles (Findable, Accessible, Interoperable, Reusable), which define the standards of research data management and are becoming a sine qua non of contemporary research projects, is also gaining in importance [8]. The FAIR principles are cross-cutting in nature: they concern both the technical accessibility and interoperability of data, discussed under data integrity, and the institutional requirements of legality and transparency, which belong to the present dimension.

Taken together, the dimensions above (epistemic distinctiveness, data integrity, documented and potential research applications, unique taxonomic status, and ethical and legal compliance) form a multidimensional profile of the scientific value of a collection, far more informative than any directly measured single figure such as the number of specimens. Such a profile makes it possible to justify investment in maintaining and developing the holdings in a defensible way. It also makes it easier to talk about their importance, both to funding bodies and to a wider public.

### 3.1 The dynamics of the scientific value of a collection

The five dimensions of scientific value described above are not fixed properties; each of them changes over time, both upwards and downwards. Understanding these dynamics is a condition of deliberate collection management: it makes it possible to identify the actions that build value and the omissions that destroy it irreversibly.

A decline in the scientific value of a collection may be material or informational. In the material dimension, the most serious threat is the physical degradation of specimens (for example desiccation, mould, pest activity or inadequate storage conditions), which renders the material useless for any future analysis. Equally serious, though less obvious, is the gradual exploitation of a collection without its replenishment: a collection intensively used for destructive research and not fed with new material gradually loses the epistemic distinctiveness that was its foundation. In the informational dimension, the key threat is the loss or degradation of metadata through the destruction of labels, the loss of field notes or the illegibility of archival records. A specimen deprived of information about the place, time and method of collection ceases to be scientific evidence and becomes merely a biological preparation of unknown provenance. A final, systemic threat is the obsolescence of the legal and ethical status of a collection: material collected without adequate permit documentation, or inconsistent with current regulations on access to genetic resources, may be excluded from scientific circulation regardless of its substantive value.

An increase in the scientific value of a collection occurs in several characteristic circumstances. First, value grows with the passage of time, when a collection documents environmental states that no longer exist (for example destroyed sites, transformed habitats, or populations that have died out naturally or been exterminated). The longer the time span of a collection and the more altered or degraded the original collecting sites, the harder comparative material is to obtain and the more irreproducible the accumulated resource becomes. Second, the value of a collection grows when changing regulations or field conditions make it impossible, or substantially harder, to obtain new material from the same localities or the same taxa. The collection then becomes the only legal source of access to a given biological resource. Third, the value of a collection grows retroactively with the development of new analytical methods: material preserved decades ago with morphology in mind suddenly becomes useful for genomics, isotope studies or metagenomics, provided it was properly stored and documented. Finally, value grows when collection material is linked to new layers of data such as georeferences, molecular sequences or high-resolution imaging, which open research avenues impossible to pursue at the time the collection was created.

These bidirectional dynamics have one fundamental practical consequence: the scientific value of a collection is the resultant of the quality of curatorial care and of research activity, not a simple function of its age, contents or size. A neglected collection loses value regardless of its potential. A collection systematically enriched and well documented gains in importance with every year, even without being supplemented with new specimens.

## 4 Incremental increase of the scientific value of natural history collections

The five dimensions of scientific value described in the previous section also define the space of possible interventions: the value of a collection can be increased systematically through purposeful curatorial and research actions that enrich existing material with new layers of information without any need to acquire new specimens. These actions differ substantially both in the resources they require and in the dimension of value they primarily strengthen, which justifies a systematic typology. As the criterion of division we adopt the object of the action, distinguishing four categories: enriching the data record, enriching the physical specimen, establishing formal status, and increasing the visibility and interoperability of the collection. The first three categories are distinguished by the object on which the curator operates: the record, the specimen and the formal status of the collection. The fourth is heterogeneous relative to the others, because it covers actions on the links between the collection and the outside world, that is, on what happens to the record and the specimen beyond the collection. We retain it as a separate category because in curatorial practice it requires different competences, different tools and a decision of its own, distinct from the other three.

The typology above is not a proposal to replace fieldwork with work on holdings already in hand. As indicated in the previous section, a collection used without replenishment gradually loses its epistemic distinctiveness, and the structural decline in the rate at which new specimens are collected is a threat in its own right to collections as research infrastructure. Incremental actions and continued acquisition of material are therefore complementary rather than alternative. What sets them apart is the ratio of effort to effect. Most of the activities described below can be carried out as part of routine curatorial work, without organising expeditions, without collecting permits and without any increase in storage space. The effect, moreover, is immediately visible in the accessibility and usability of the data. This makes them a rational starting point for increasing the value of a collection, particularly where funds for fieldwork are limited.

The four categories are not, however, equivalent in terms of the order in which they should be implemented. Enriching the data record is the natural first step, because it requires neither instruments nor permits nor access to the original, and its effects are fully reversible, so the risk of a wrong decision is smallest. Enriching the physical specimen already requires laboratory infrastructure, and some of the operations are irreversible, so it is sensible to undertake them where the record is described well enough for the result of an analysis to be assigned unambiguously to a specimen and a site. Establishing formal status is largely independent of the two preceding categories, but it presupposes taxonomic competence and access to the legal documentation of the holdings. Increasing visibility comes last, because publication in data aggregators makes sense only after the record has been standardised: a collection made available with incomplete or inconsistent metadata reduces trust in it instead of building it. This order is not a rigid schedule but a default rule for holdings in which none of the dimensions is yet being developed systematically.

### 4.1 Enriching the data record

Actions in this category operate on the description of the specimen without any intervention in the physical material, which makes them the most accessible and most cost-effective way of increasing the scientific value of existing holdings. Their common denominator is the completion of missing information, or the conversion of information hitherto hidden or hard to reach in labels, field notes and old catalogues, into structured, machine-readable data integrated with the specimen record.

Fundamental here is the retrospective georeferencing of historical labels, that is, the conversion of descriptive locality data into WGS84 geographic coordinates with an assigned measure of spatial uncertainty [24]. Georeferencing opens the possibility of including historical specimens in analyses of species distributions, in ecological niche modelling and in studies of range change, which without such data would be impossible or seriously affected by sampling bias [25]. A complementary step is the retroactive attachment to records of environmental data corresponding to the dates and localities of collection (for example climatic values from the WorldClim, CHELSA or ERA5 databases), which allows correlations between organismal traits and environmental conditions to be analysed without any additional field measurements [26].

Standardising metadata to the Darwin Core or ABCD schemas allows data to be aggregated directly with other collections without costly conversion and is a precondition of publication in global aggregators [7]. Even a minimal extension of the record to include the collecting method, habitat type, phenological stage and determination history significantly raises the usefulness of the data in comparative studies. Revision of taxonomic identifications, documented as a separate entry in the determination history of the catalogue record, improves the taxonomic quality of a collection without any intervention in the specimens and provides the basis for their reliable inclusion in systematic syntheses. Attaching to records the citations of papers in which a given specimen served as a reference, with the DOI and the type of use, creates a documented trajectory of the scientific life of the specimen and secures knowledge that would otherwise remain only in the memory of individual researchers. Actions in this category affect above all the integrity and accessibility of data and the documented and potential research applications of the collection [6].

### 4.2 Enriching the physical specimen

Actions in this category involve direct intervention in the biological material and aim to extract from the specimen information unavailable at the level of the label or catalogue record alone. Foremost among them is the taking and cryopreservation of tissue vouchers, that is, small samples taken at the moment the specimen is prepared and stored at −80°C or in RNA-stabilising solutions [27]. They secure genetic material for analytical methods that were outside the standard laboratory repertoire at the time of collection, such as comparative genomics, transcriptomics and proteomics. The window for taking tissue cannot be opened retrospectively, which makes this operation a particularly important element of routine collecting practice.

A complementary approach is DNA barcoding [9], the sequencing of standard genetic markers (COI for animals, ITS or rbcL for plants and fungi) combined with the deposition of sequences in the BOLD and GenBank databases with an explicit link to the catalogue record of the specimen [10]. This transforms a physical specimen into a node of the global molecular network and is one of the most effective ways of increasing its citability. For historical museum specimens, the retroactive recovery of DNA using modern amplification methods makes it possible to obtain sequences even from preparations more than a century old, opening access to phylogeographic and evolutionary data on a time scale inaccessible to field research [4], [28].

High-resolution imaging, both photography under standardised conditions (with a colour chart and constant lighting) and more advanced techniques such as scanning electron microscopy (SEM) and computed microtomography (micro-CT) [29], creates durable digital representations of the specimen that are available remotely and protect the original from excessive handling. Micro-CT data can be deposited in repositories such as MorphoSource, where they become a publicly available morphological resource independent of the physical location of the specimen [30]. Systematic morphometrics, comprising the measurement of a standardised set of characters for large series of specimens and the publication of raw data in repositories such as Dryad or Zenodo, gives a collection comparative value for future meta-analyses that could not be obtained from verbal descriptions alone [31]. Actions in this category require access to the original material, often specialised infrastructure, and in some cases consent to destructive sampling. Their effect is primarily to strengthen the epistemic distinctiveness of the collection, understood as its capacity to supply data not replicable from other sources [19].

### 4.3 Establishing formal status

Actions in this category are permanent, and their effects reach beyond the individual collection and enter the formal apparatus of science. They comprise nomenclatural acts, registration in international databases and verification of the ethical and legal compliance of the holdings. The starting point should be a systematic inventory of the collection with respect to type specimens: holotypes, syntypes, lectotypes, neotypes and paratypes, whose status is sometimes undocumented or dispersed in older literature [12]. Every type so established should be registered in ZooBank (for animals) or in IPNI and Index Fungorum (for plants and fungi) with an assigned persistent LSID [13]. Where the original description of a species rested on a series of syntypes without the designation of a holotype, the designation of a lectotype from existing collection material is a formal nomenclatural act that can be published as a short taxonomic paper [12]. A single such designation may establish the status of a collection as a reference collection for decades. In parallel, it is necessary to verify collecting permits, to document the policy on destructive sampling and to register material in accordance with the requirements of the Convention on Biological Diversity (CBD) and the Nagoya Protocol [32]. In practice, legal documentation conditions the circulation of material as much as its substantive quality: an international loan and publication in a peer-reviewed journal today require both. Actions in this category strengthen above all the unique taxonomic status and the ethical and legal compliance of the collection [22], [23].

### 4.4 Increasing visibility and interoperability

Actions in this category do not change the contents of a collection but determine the extent to which it is genuinely accessible and useful to the wider scientific community. A collection without digital visibility remains a local resource regardless of its substantive value. Publication of data in GBIF, for example through the Integrated Publishing Toolkit (IPT), gives every data set global visibility and a persistent DOI [33]. This makes it possible to transform a local resource into a node of the global infrastructure of biodiversity knowledge. Depositing DNA sequences in the BOLD and GenBank databases with an explicit link to the specimen record incorporates the collection into a molecular network cited by a growing number of systematic and ecological studies [9], [10]. Linking records to TreatmentBank and to the CrossRef system creates a mutually verifiable network of specimen-publication links, making it possible to track the actual scientific use of a collection, which in turn may provide the basis for its fair assessment by funding bodies [34].

The architecture of the future for a full implementation of these principles is the developing Extended Specimen standard (DiSSCo/iDigBio) [14], which allows a specimen record to be extended with any linked data objects (a DNA sequence, an SEM image, climatic data, the results of isotope analyses) as attributes of a single identified digital object. This standard aligns directly with the FAIR principles (Findable, Accessible, Interoperable, Reusable) [8], which are becoming a requirement of publicly funded research projects. Actions in this category affect almost all the dimensions of scientific value simultaneously, and their effects accumulate over time as the number of users of global data aggregators grows [6].

The diagram below (Figure 1) presents the structure as a bipartite graph. Solid lines (colour-coded) indicate the principal, direct influence of a given category of actions on a specific dimension of value: the physical specimen (yellow) feeds primarily epistemic distinctiveness, the data record (blue) feeds integrity and applications, and formal status (red) feeds taxonomic status and compliance. Dashed lines (grey) show the cross-cutting character of visibility, a category that strengthens all five dimensions without any one of them being its exclusive target. The links shown are dominant rather than exclusive. The standardisation of metadata, assigned to enriching the data record, also raises epistemic distinctiveness, because it makes series of specimens comparable across time and between sites. Second-order dependencies have been omitted for the legibility of the diagram. The diagram presents the static structure of the framework, whereas the dynamics of value discussed earlier remain a separate perspective.

**Figure 1.**
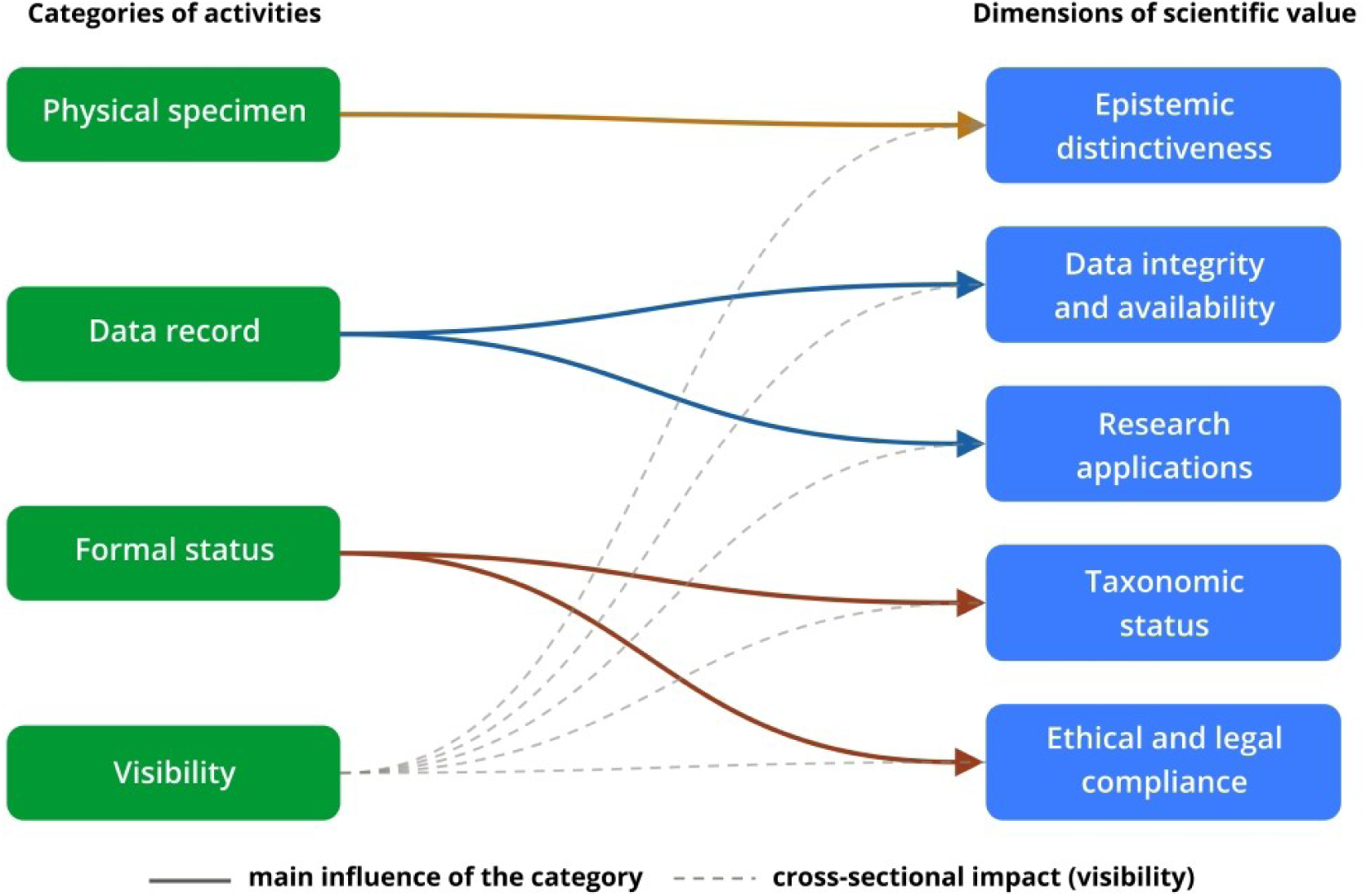
Categories of actions and the dimensions of the scientific value of a natural history collection

The framework from Section 3 and the typology from the present section remain, so far, theoretical constructs. In the next section we test them against the material of two collections held in the Natural History Collections of Adam Mickiewicz University, showing which of the actions described have actually been carried out in them, which dimensions of value this has raised, and what remains to be done.

## 5 The framework applied: two collections of Adam Mickiewicz University in Poznań

Among the disciplines that use natural history collections, faunistics is the one in which dependence on accumulated material is most direct. The composition and distribution of fauna in a given area can be reconstructed only from voucher specimens and the metadata associated with them. Every conclusion about a change of range, about a species disappearing from a site or about the appearance of a new species requires comparative material from an earlier period. For this reason the objection sometimes raised, that faunistics is merely an inventory activity rather than science, is hard to sustain. It is precisely faunistic time series that make it possible to assess the conservation status of threatened species, to monitor environmental change and to take decisions on the sustainable management of natural resources. The collections discussed below provide the basis for research of exactly this kind, and at the same time allow the successive dimensions of collection value distinguished in Section 3 to be traced.

Abstract criteria of scientific value acquire their full meaning only in confrontation with concrete research material. Below we illustrate them with selected collections of the Natural History Collections. This unit, museum in character, was established in 2004 at the Faculty of Biology of Adam Mickiewicz University in Poznań, and its task is to assemble, properly store and make available scientific natural history collections comprising zoological holdings, herbaria and osteological collections. The Natural History Collections were created as a place to house collections considerably older than the unit itself, assembled in the departments of the Faculty since the mid-twentieth century, which explains the discrepancy between the date the unit was founded and the dates at which the two collections discussed below began.

### 5.1 First example: the collection of soil samples and samples from various microhabitats

#### Origin of the collection

The soil samples held are the product both of the research activity of the group of acarologists initiated by Professor Jan Rafalski and continued to this day by his students, and of students active in the Invertebrate Research Section of the Naturalists’ Student Society, of participants in research expeditions, and of people entirely unconnected with science who supplied samples with no expectation of return. In this way, from the establishment of the Department of Animal Morphology in 1961, close to 40,000 samples have been assembled, not only from Poland but also from many countries on all continents. A detailed characterisation of the collection and the history of its formation are presented in the first part of the Catalogue of soil samples [35].

#### Basic information on the holdings from the territory of Poland

Of the nearly 40,000 samples held in the collection, 7,459 soil samples and samples from various ephemeral microhabitats have so far been sorted and fully catalogued [35,36,37]. The material came from three basic habitat types: open non-forest habitats, forests and scrub, and various ephemeral microhabitats (merocoenoses), from the whole territory of Poland and from the years 1931-2025 (the period of sample collection). Open environments are represented by 10 types, of which meadows and rock swards were sampled most frequently. Forests are represented by 13 communities, with the greatest number of samples taken in oak-hornbeam forests, beech forests, riparian forests, pine and spruce forests, and parks. The merocoenoses comprise 10 microhabitats, of which dead wood is the most abundantly represented, followed by tree hollows, the nests of birds and small mammals, and ant nests. A substantial part of the samples comes from protected areas: 20 national parks and 200 nature reserves, and in some sites the samples were taken before legal protection was introduced. The collection covers the full altitudinal range of the country, from depressions to the highest peaks of the Tatra Mountains. In some sites, such as the oak-hornbeam reserves of western Wielkopolska and the Cisy Staropolskie im. L. Wyczółkowskiego reserve at Wierzchlas, samples have been taken regularly for many years. An exceptionally long study period characterises the Jakubowo reserve, where soil fauna has been monitored without interruption since 1973, giving more than fifty years of comparable samples from a single site. This temporal continuity and breadth of habitat coverage are the direct basis of the epistemic distinctiveness of the collection in the sense adopted in Section 3. The distribution of the sampling localities of the catalogued samples is shown in Figure 2.

**Figure 2.**
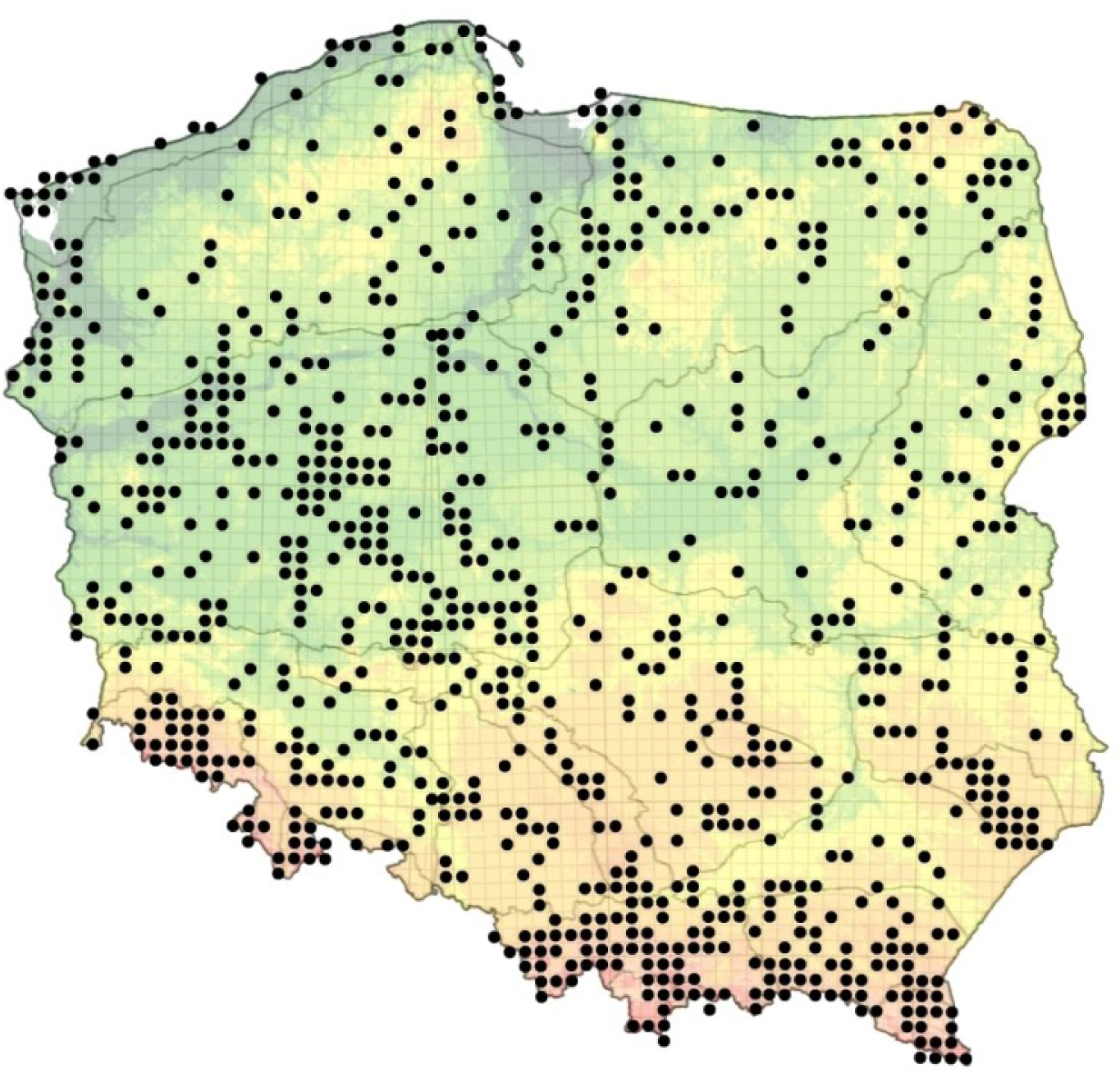
Location of the sampling sites of the 7,459 samples in Poland (in the 10×10 km UTM system)

Cataloguing here means the full description of a sample in the database, including locality, date, collector, habitat type and extraction method, and it is an operation distinct from examining a sample for a particular invertebrate group. The numbers of samples examined in studies of certain groups of oribatid mites (Oribatida) or of mites of the suborder Uropodina, given later in the text, refer to this second operation and also include samples whose description in the database remains incomplete. These two numerical orders neither add up nor contain one another.

#### Methods of collection and extraction

Samples in the collection are either qualitative or quantitative. The former are sieved samples of litter and soil, or unsieved samples from various ephemeral microhabitats. Quantitative samples were taken in the field by a volumetrically standardised method (cylinders of defined diameter and depth of insertion), which ensures comparability of results between sites and between years. The material was extracted in the laboratory by the thermal method using a Berlese-Tullgren funnel, in which illumination and the gradual heating of the sample from above force the invertebrates to migrate downwards into a vessel with preserving fluid. This method is the standard in soil fauna research, particularly for mites and springtails, and allows these invertebrates to be extracted efficiently without destroying delicate specimens. Samples from moist habitats and from habitats with a high organic matter content were additionally processed by flotation and wet sieving. The material obtained was preserved in 75% ethanol, which permits long-term storage both of specimens intended for morphological study and of genetic material. In the latter case the specimens are additionally kept at low temperatures.

Changes in the way samples are stored (Figure 3) followed the sequence below. Glass Weck jars (1), in which samples were kept from the beginning of the collection, were replaced in the late 1990s by plastic containers with screw caps and site labels (2). In both these solutions individual samples were kept in glass tubes plugged with cotton wool, which required constant topping up of alcohol, and the jars and containers themselves took up a great deal of space. Access to individual samples was also difficult, since several or a dozen had to be looked through before the right one was found. Today each sample goes into a separate plastic tube with a sealed cap and its own label, in two sizes chosen according to the volume of material (3). The tubes are placed upright in partitioned boxes (4, 5). This gives direct access to every sample without removing the others and markedly reduces the space occupied.

**Figure 3.**
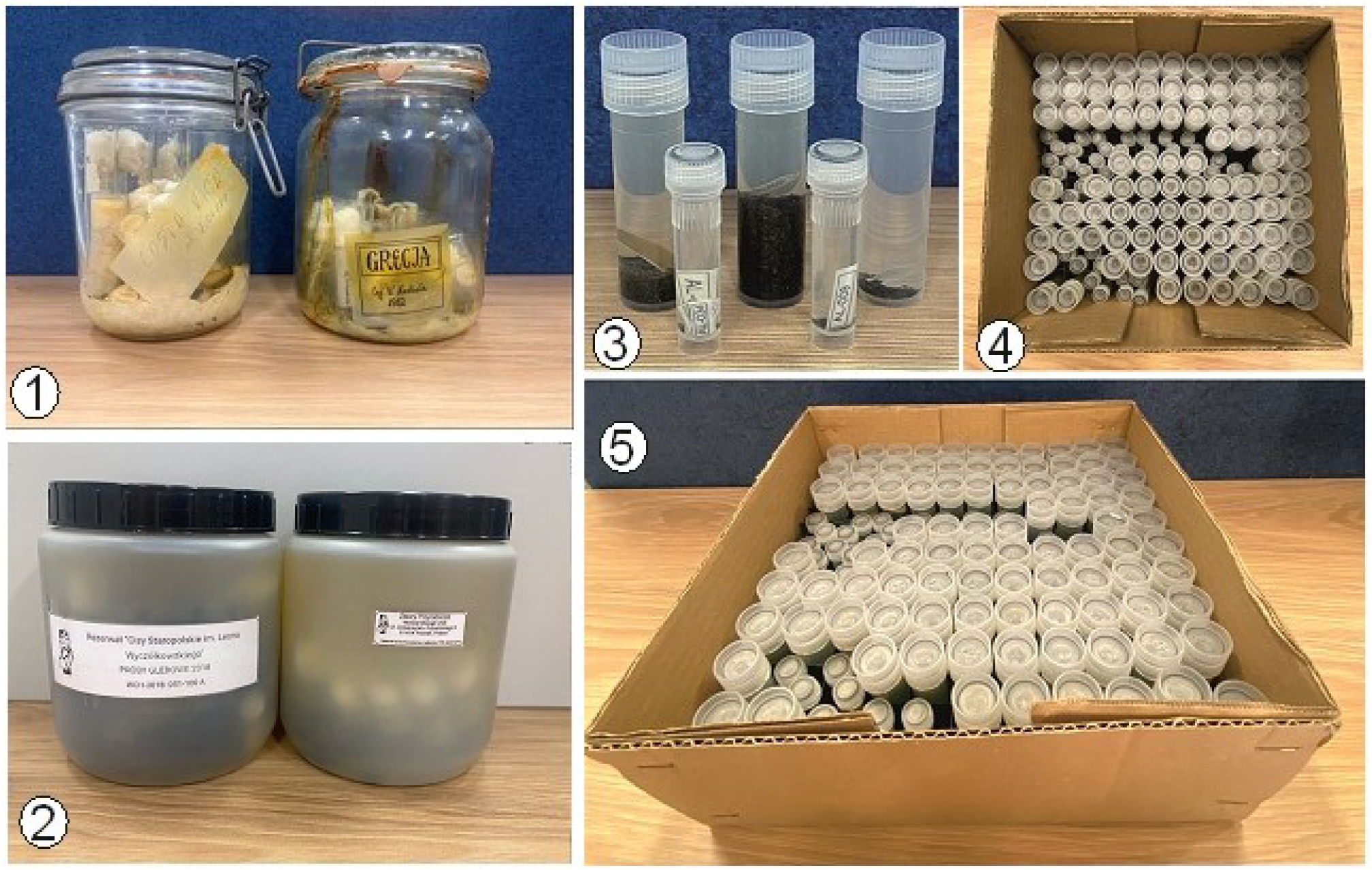
Evolution of sample storage: 1 - glass Weck jars (twentieth century), 2 - plastic containers with screw caps (turn of the twentieth and twenty-first centuries), 3 - plastic tubes with sealed caps in two sizes, 4 and 5 - tubes arranged in partitioned boxes (present day)

#### Methods of study: preparation, microscopy and molecular methods

Morphological study of the specimens was carried out by light and phase-contrast microscopy after clearing in lactic acid or potassium hydroxide (KOH) and mounting in Hoyer’s medium or glycerine. The microscope slides so produced allow identifications to be revised repeatedly. Their durability is not, however, unlimited: Hoyer’s medium crystallises and dries at the edges over time, which requires the slides to be ringed and the state of the collection to be checked periodically. This is a concrete example of the material degradation of value described in Section 3.1. Abandoning such checks would mean losing not the specimen itself but the possibility of verifying its identification, and hence the evidential character of the whole series. For selected specimens, particularly in the descriptions of new species, scanning electron microscopy (SEM) was used, allowing the microstructure of the cuticle, the chaetotaxy and the gnathosoma to be imaged at a resolution unattainable by light methods. For the two leading groups (Oribatida and Uropodina) more than 5,000 SEM images have been produced, which have fed the iconographic collection of the Natural History Collections and constitute a durable visual resource available for future taxonomic revisions. This is a direct realisation of actions from the category of enriching the physical specimen discussed in Section 4. In the area of molecular methods, DNA barcoding was carried out for more than 100 species of Uropodina and for selected representatives of Oribatida, sequencing standard genetic markers and depositing the sequences obtained in global gene banks (GenBank, BOLD). Molecular analyses initially required destructive sampling of tissue from individual specimens. Being irreversible, this is possible only at the moment the material is being processed, which underlines the importance of the routine securing of tissue discussed in Section 4. Extraction of genetic material now allows the cleared specimens to be preserved, which makes it possible to examine them under a light microscope and thus guarantees correct assignment to species. The results of these analyses have transformed physical specimens into nodes of the global molecular network, greatly increasing their citability [38], [39].

#### The digital system for managing collection data

The transition from analogue to digital recording of the soil sample collection proceeded gradually (Figure 4). The spread of personal computers in the early 1980s made it possible to convert the descriptions of samples and of their contents into digital form.

**Figure 4.**
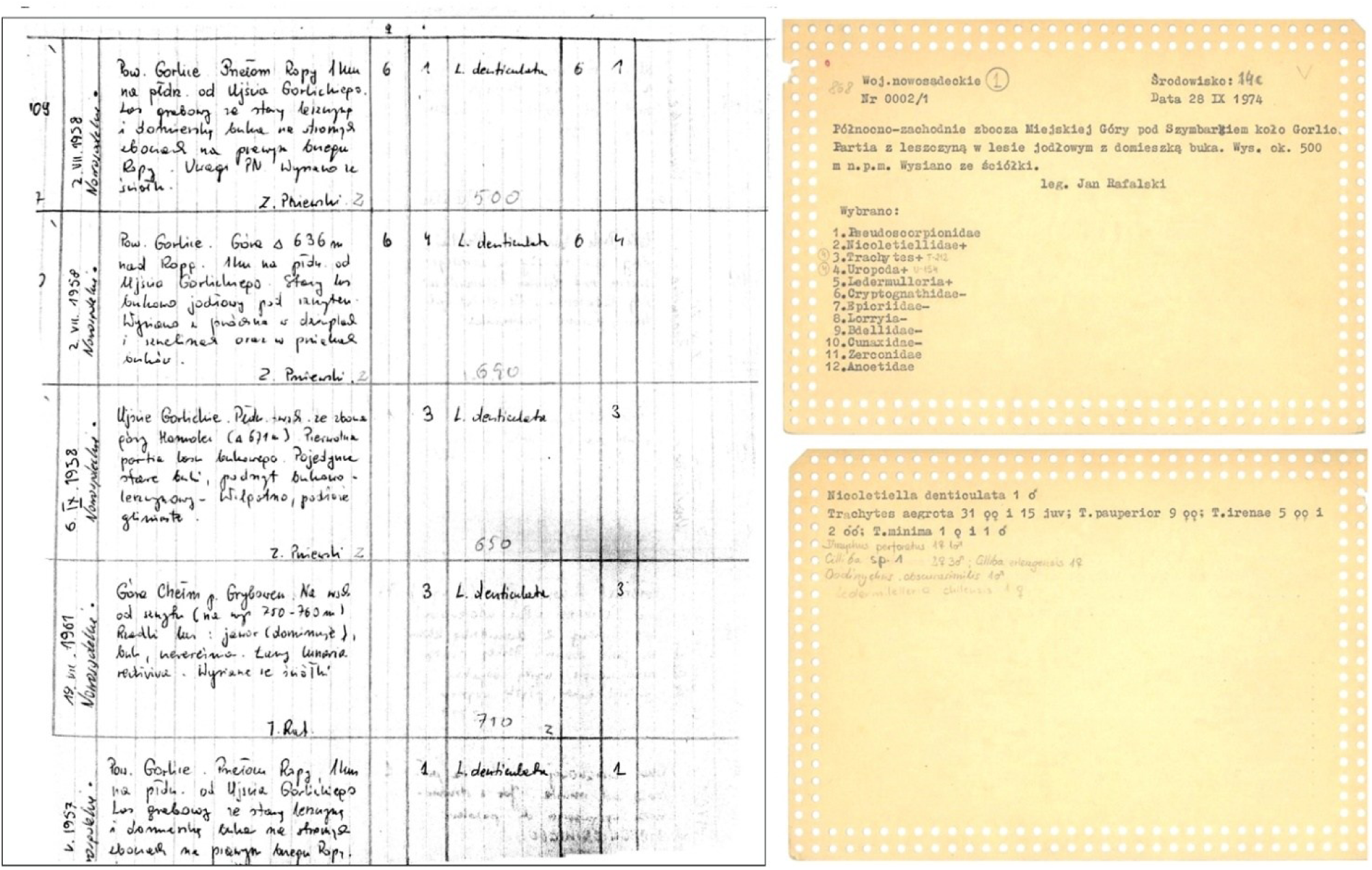
Evolution of analogue recording: from conventional notebook entries to punched cards

A key element of the collection infrastructure is the ANALIZATOR program, developed by the Desmodus company and in use since 1999. It functions as a digital database, an extended catalogue of sample contents, integrated with a metadata analysis module. The system allows samples to be searched rapidly by locality, date, collector, habitat type and taxonomic composition, which in practice transforms the physical collection into a data structure fit for research queries. The full range of metadata has so far been achieved for 7,459 samples; the remainder appear in the database at a basic level, which limits the possibility of querying certain fields. The metadata of a sample comprise: a verbal locality with the possibility of georeferencing, the date of collection, the name of the collector, habitat type, altitude above sea level, extraction method, and a taxonomic composition that is filled in progressively. This corresponds to the data record standards discussed in Section 4, although full standardisation to the Darwin Core schema remains an action for the future.

#### Use made of the collection to date

A substantial part of the samples was assembled as research material for degree theses carried out by staff of several departments of the Faculty of Biology. Invertebrate groups selected from these samples became the basis of doctoral and habilitation dissertations, most of which were published as monographs. It is worth noting that each member of staff working on a given group of mites drew equally on their own samples and on samples collected by others. In acarology alone, 9 doctoral dissertations and 3 habilitations have been based on the sample collection (Table 2).

**Table 1.** presents the same material as a checklist, of use when planning work on a specific collection. Categories of incremental actions: example operations, the dimensions of scientific value they strengthen, and the resources required

| Category of actions | Example actions | Dimensions strengthened directly | Resources and conditions required |
| --- | --- | --- | --- |
| Enriching the data record | georeferencing of historical labels; attaching environmental data to the record; standardisation to Darwin Core or ABCD; revision of identifications with a determination history; recording of specimen citations | data integrity and accessibility; documented and potential research applications | curatorial time and access to the database; no intervention in the material; lowest barrier to entry |
| Enriching the physical specimen | taking and cryopreservation of tissue vouchers; DNA barcoding; retroactive recovery of aDNA; SEM and micro-CT imaging; morphometrics of large series | epistemic distinctiveness; documented and potential research applications | access to the original; specialised instrumentation; consent to destructive sampling; some actions irreversible |
| Establishing formal status | inventory of type specimens; registration in ZooBank, IPNI or Index Fungorum; designation of lectotypes; verification of permits and of Nagoya Protocol compliance | unique taxonomic status; compliance with ethical and legal standards | taxonomic competence; access to the legal documentation of the holdings; permanent effect, reaching beyond the collection |
| Increasing visibility and interoperability | publication in GBIF via IPT; deposition of sequences in BOLD and GenBank; linking to TreatmentBank and CrossRef; implementation of Extended Specimen | all five dimensions, none exclusively | prior standardisation of the record; IT support; effect accumulates over time |

**Table 2.**
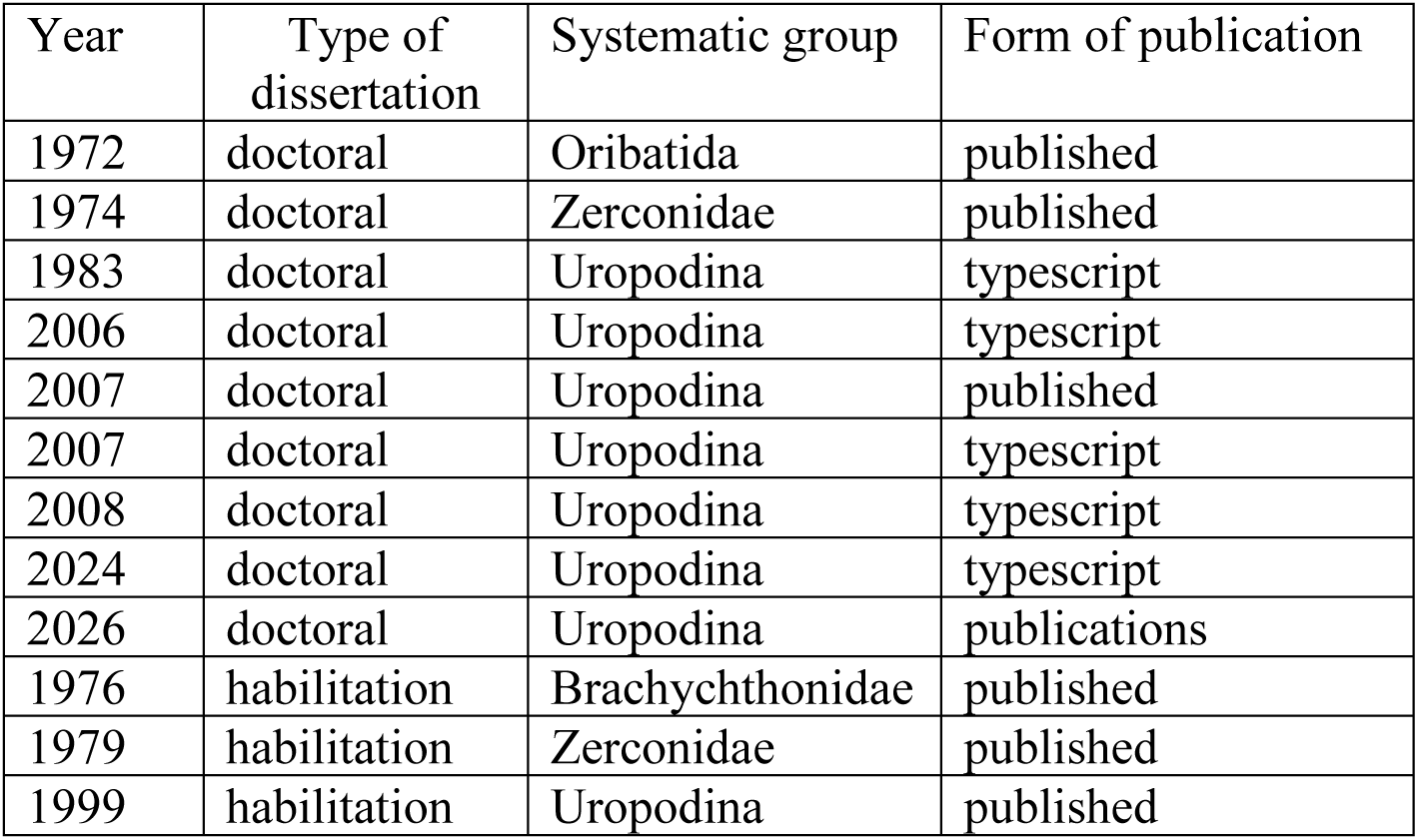
Doctoral and habilitation dissertations based on the collection of soil samples, by year, systematic group and form of publication. An en dash marks an entry for which the year could not be established from the records held.

An even more numerous group are students using the samples, or the metadata already available, for their diploma theses (bachelor’s and master’s). Some of them draw exclusively on samples already collected, while others base their theses on material they have gathered themselves, which after processing feeds the collection. The number of completed bachelor’s and master’s theses based on the collection of soil samples has passed 100.

The existence of the collection of soil samples made it possible to produce monographic treatments of several groups of mites on a national scale, something achieved so far in only a few countries [40,41,42,43,44,45,46]. This scale would not have been attainable without the collection as shared infrastructure, in which each researcher drew in parallel on their own samples and on those of others, a model instance of the synergistic effect described in Section 3 as a feature of epistemic distinctiveness. Comparable treatments of some groups of mites on a national scale have been produced outside Poland, in Slovakia among other countries [47]. The total number of publications based on material from this collection, both Polish and exotic, runs into the thousands; this is an estimate derived from the publication output of the Poznań acarologists. No systematic bibliometric survey has yet been carried out.

#### Benefits of holding the collection

Access to the sample collection offers researchers and students a range of possibilities and is a substantial aid to research work. First, it allows the use of material whose independent collection would far exceed the capacity of a single person, which saves time and money. Second, it opens access to historical material, often gathered in places that have since been profoundly transformed or have ceased to exist altogether (felled woodland, the beds of artificial water bodies, built-up areas). Third, samples taken at different periods at the same site, whether in a national park, a nature reserve or any other locality, make it possible to track changes in the species composition and structure of invertebrate assemblages. Finally, the exotic samples give access to material from many countries on different continents: for a taxonomist this means the possibility of obtaining specimens from all over the world without organising costly and often dangerous expeditions.

The samples collected contain various groups of soil fauna. Among them are specimens of extremely rare species that are difficult to obtain in the field. An example is the small, heavily sclerotised myriapods of the genus Brachypauropus Latzel, 1884: three species representing this group have so far been recorded, after examining more than 20,000 samples, on only two occasions, in a total of six individuals [48] (Błoszyk 2025, unpublished data). Taken together, across all the groups of mites studied so far, the samples held in the collection have yielded more than 1,000 species new to science and several hundred species new to the fauna of Poland.

#### Derivatives of the soil sample collection as added value

One of the effects of the scientific use of the soil sample collection is the secondary creation of collections of the invertebrate groups studied. We take as our example the two best-studied groups of mites, namely the oribatid mites of the group Ptychoidea (Oribatida) and the mites of the suborder Uropodina (Mesostigmata). For the former, more than 800 species new to science from all over the world have been described on the basis of the soil sample collection assembled over the years, in this group alone. This gave rise to a collection of nomenclatural types containing the holotypes and paratypes of these species (Figure 5).

**Figure 5.**
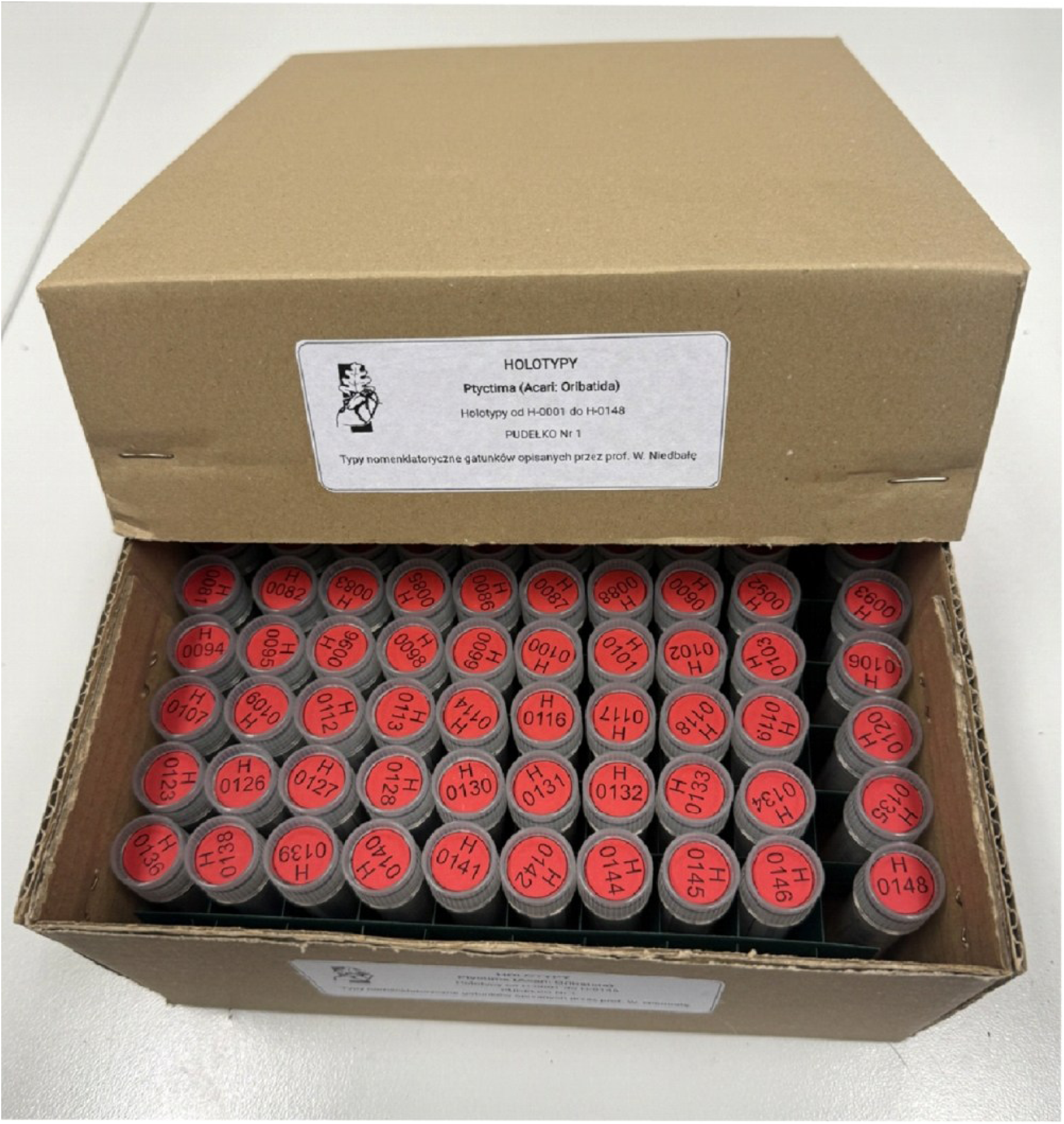
The collection of nomenclatural types (holotypes) of Ptyctima

In addition, 44 species of this group of mites were recorded in 2,500 samples collected across Poland, in a total of more than 61,000 specimens, which have been preserved as a further collection. The results of molecular studies based on these specimens have enriched the holdings of the gene bank, and their SEM images have enriched the iconographic collection of the Natural History Collections. For the Uropodina, fewer species new to science have been described, but more than 200 from various geographic regions of the world still await description. From Poland, 150 species were recorded in nearly 25,000 samples, in a total of about 500,000 specimens. These specimens, kept in fluid or as microscope slides, now constitute a separate collection deposited in the Natural History Collections at the Faculty of Biology, AMU, in Poznań. As in the case of the oribatid mites of the group Ptychoidea (Oribatida), molecular analyses were carried out for more than 100 species, and some of these data have enriched global gene banks. Several thousand SEM images produced from this collection have also entered the iconographic collection of the Natural History Collections.

Taken together, the two best-studied groups convey the scale of the whole undertaking: 27,500 samples examined, close to 200 species recorded in Poland, and more than 560,000 identified specimens preserved as separate collections. Set against the nearly 40,000 samples that make up the entire holding, these figures also show how large a part of the material remains unexamined for the other groups of soil fauna.

The trajectory described, from a raw soil sample through morphological preparations and SEM images to DNA sequences in gene banks and collections of nomenclatural types, is an empirical illustration of the incremental increase of the scientific value of a collection in all four categories of action distinguished in Section 4. None of these steps required a repeat field trip; the derivative collections arose solely from material already deposited. The base collection itself, however, continues to be fed with new samples, which, in line with the qualification made in Section 4, remains a condition of maintaining its epistemic distinctiveness in the longer term.

#### Untapped potential

Despite the intensive use made of it so far, the collection of soil samples holds substantial research potential that can be activated at any time. The samples stored in ethanol are a resource for analyses of environmental DNA (eDNA) and of soil metagenomics. The 75% concentration used is a compromise between the requirements of morphological preparation and the preservation of nucleic acids, which realistically limits such analyses to short barcode amplicons and metabarcoding rather than to the sequencing of long genomic fragments. The scale and temporal span of the collection nevertheless mean that even so limited a range provides comparative material unobtainable in any other way. This is a second instance of the dynamics described in Section 3.1. A preparation decision taken decades ago, entirely rational from the standpoint of morphology, today limits the range of molecular analyses. Part of the potential was therefore lost before the methods that could have exploited it even existed. Those methods were not part of the standard laboratory repertoire when most of the samples were being collected. Retrospective georeferencing of samples described by verbal locality would open the possibility of including historical material in global analyses of distribution and in models of climatic change. Publication of the catalogue data in GBIF would give the collection global visibility and a persistent DOI, transforming a local resource into a node of the international infrastructure of biodiversity knowledge. Publications in GBIF are planned as the next stage of work on the collection and will be carried out progressively, together with the successive printed volumes of the Catalogue of soil samples. An intermediate step that has to be taken first is the mapping of the ANALIZATOR database structure onto the Darwin Core schema, above all the completion of records with geographic coordinates and the harmonisation of the vocabularies of habitats and collecting methods. Only after this standardisation will the collection data be exportable as a Darwin Core Archive and publishable through the Integrated Publishing Toolkit.

### 5.2 Second example: the conchological collection of shells and of field observation data of the Roman snail (Helix pomatia) in Poland

#### Origin of the collection

The collection dates back to the 1960s. It was created in the Department of General Zoology on the initiative of Professor Jarosław Urbański. Initially the holdings comprised shells and alcohol-preserved snail bodies from various regions of Poland, collected by members of the department. Gradually they were also supplemented with voucher material from master’s theses (Figure 6). At the beginning of the twenty-first century the collection numbered more than 30,000 catalogued Roman snail shells with precise localities of origin and data on the environment of occurrence, together with a dozen or so Weck jars holding preserved bodies. A rapid expansion of the collection, accompanied by a fundamental change in its character, began in 2009. The collecting of shells was replaced by field observation using GPS devices, which made it possible to record the location of individual specimens directly in the field and then, using GIS systems, to visualise their distribution. Extensive surveys of the Roman snail commissioned by the Regional Directorates for Environmental Protection in several provinces (Podlaskie, Wielkopolskie, Lubuskie, Kujawsko-Pomorskie, Dolnośląskie and Zachodniopomorskie) enriched the collection with metadata derived from field observation.

**Figure 6.**
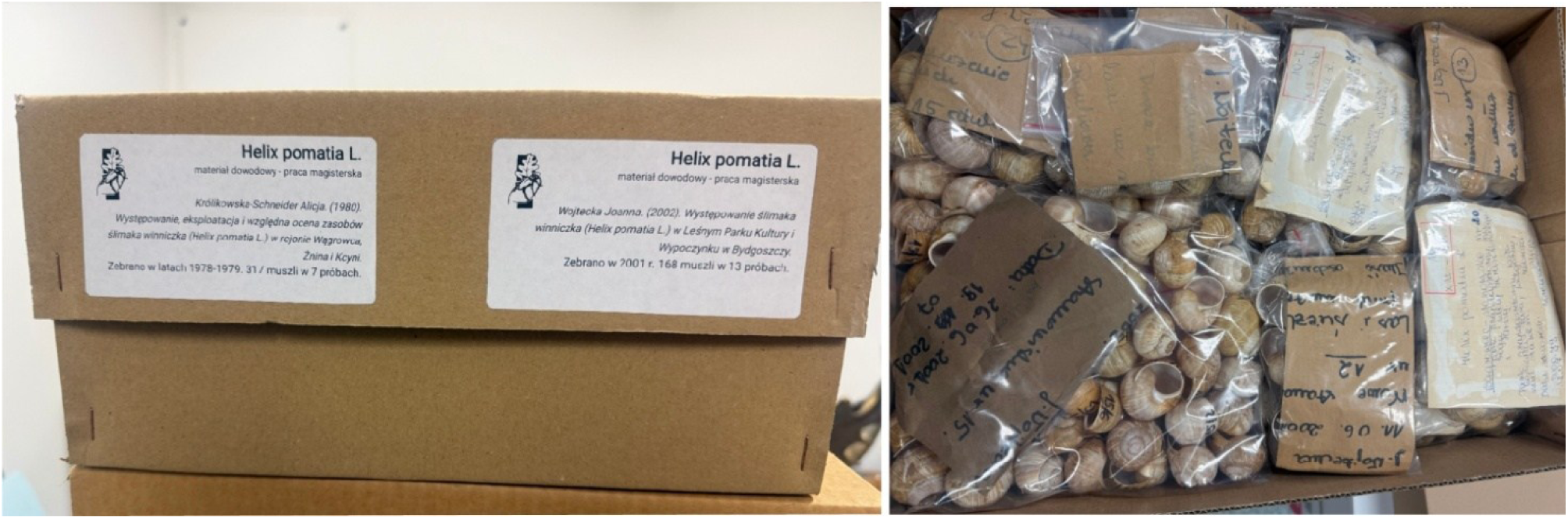
The conchological collection: voucher material for a master’s thesis

#### Basic information on the holdings

The core of the collection still consists of shells from sites scattered across the country, provided with a description of the place and date of collection and a characterisation of the habitat, supplemented by alcohol-preserved snail bodies. Each series is accompanied by biometric measurements of the shell, its diameter and height, and by the weight of the snail. Shell diameter is the feature on which the statutory criteria for permissible commercial harvesting of the snail are based. A second and considerably younger layer of the collection consists of the field observation records mentioned above: the positions of individual specimens recorded with a GPS receiver together with a description of the site and, for part of the material, with an individual mark on the specimen allowing it to be found again in later seasons (Figure 7). This second layer has no material counterpart; its only carrier is the record in the database, which makes it particularly susceptible to the loss of value through metadata degradation described in Section 3.

**Figure 7.**
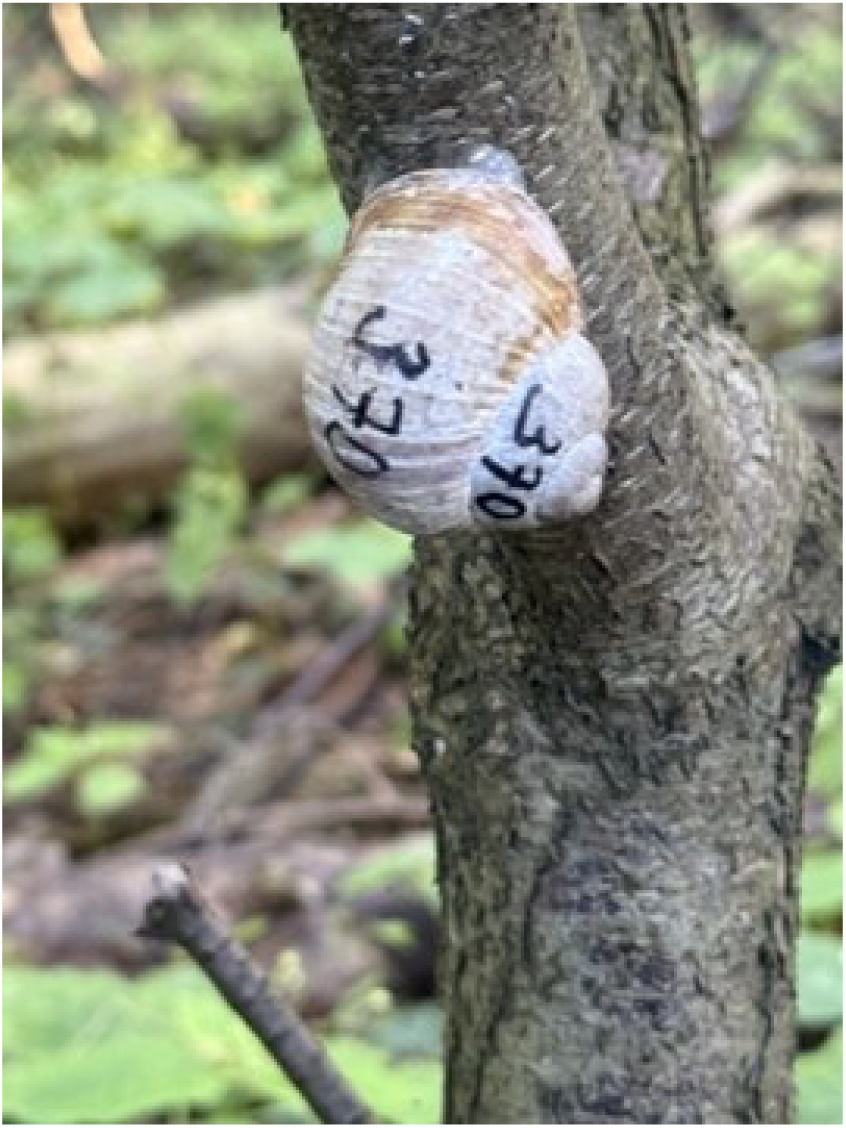
A marked snail specimen used in metapopulation studies

The observation layer is made up of data from 13 field campaigns carried out between 2009 and 2024 in six provinces (Kujawsko-Pomorskie, Wielkopolskie, Lubuskie, Podlaskie, Zachodniopomorskie and Dolnośląskie), commissioned by the Regional Directorates for Environmental Protection. In the course of these campaigns 4,563 site visits were made, the Roman snail was confirmed at 3,041 of them, which gives an overall frequency of occurrence of 66.6%, and more than 128,000 individuals and shells were recorded. Four provinces were surveyed more than once, Kujawsko-Pomorskie five times, which yields comparative series from the same areas spanning more than a decade. Table 3 summarises these campaigns by province and year. It documents the growth of the observation layer rather than the biology of the species: what matters here is the number of records accumulated, the geographic coverage and, above all, the fact that four provinces were surveyed repeatedly, which yields comparative series that cannot be obtained retrospectively. The physical part of the collection, accumulated mainly before 2009, comprises the catalogued shells and the alcohol-preserved bodies described above.

**Table 3.** Growth of the observation layer of the Roman snail collection: field surveys carried out in 2009–2024 in six provinces of Poland, commissioned by the Regional Directorates for Environmental Protection. Records are all occurrences of live individuals and empty shells registered during a campaign. The report of the 2016 survey in Dolnośląskie is not available, so its records are absent from the total.

| Province | Year | Sites checked | Sites with snails | Frequency | Records |
| --- | --- | --- | --- | --- | --- |
| Kujawsko-Pomorskie | 2009 | 281 | 176 | 63% | 7,213 |
| Kujawsko-Pomorskie | 2010 | 334 | 218 | 65% | 3,635 |
| Kujawsko-Pomorskie | 2012 | 464 | 217 | 47% | 5,395 |
| Kujawsko-Pomorskie | 2017 | 104 | 101 | 97% | 5,335 |
| Kujawsko-Pomorskie | 2018 | 299 | 210 | 72% | 14,364 |
| Podlaskie | 2011 | 314 | 181 | 57% | 4,241 |
| Lubuskie | 2012 | 325 | 192 | 59% | 2,924 |
| Lubuskie | 2016 | 203 | 188 | 93% | 4,342 |
| Dolnośląskie | 2016 | 634 | 380 | 59% | no report |
| Wielkopolskie | 2015 | 667 | 460 | 69% | 51,440 |
| Wielkopolskie | 2020 | 500 | 351 | 70% | 15,118 |
| Zachodniopomorskie | 2023 | 219 | 182 | 83% | 4,367 |
| Zachodniopomorskie | 2024 | 219 | 185 | 84% | 9,800 |
| **Total** | **2009–2024** | **4,563** | **3,041** | **66.6%** | **128,174** |

#### Methods of collection and documentation

The material was obtained by hand collecting on designated sample plots, with fieldwork concentrated in the period of spring activity of the snails, which coincides with the statutory window for commercial harvesting, that is, from 20 April to 31 May. For each individual, shell measurements were recorded that allow it to be assigned to a size class. This has direct practical significance. The Regulation of the Minister of the Environment of 16 December 2016 on the species protection of animals permits the collection only of individuals with a shell diameter of not less than 30 mm, and in the Opolskie, Śląskie, Małopolskie, Świętokrzyskie, Podkarpackie and Lubelskie provinces of not less than 31 mm. The size structure of a population therefore translates directly into whether, and on what scale, harvesting in a given area may be permitted. After processing, the shells were dried and catalogued together with a site label, and the bodies were preserved in alcohol. Since the switch to satellite-based documentation, the physical holdings have been supplemented selectively, to the extent needed to verify measurements, and the principal product of fieldwork has become the observation record.

#### The digital system for managing collection data

Recording positions in geographic coordinates transformed the collection into a layer of spatial information that can be combined with any other thematic layers: land cover, the boundaries of municipalities and provinces, protected areas or climatic data. A single record comprises coordinates, a date, a habitat description and, where possible, the identifier of a marked individual and the results of measurements. This structure corresponds to the core of the Darwin Core schema, which means that the collection can be made available in data aggregators without rebuilding the data model, simply by mapping field names. This corresponds to the actions from the category of enriching the data record discussed in Section 4.

#### Use made of the collection to date

For decades, information derived from the metadata of the collection provided the basis for setting limits on the commercial harvesting of the Roman snail in Poland, initially in decisions of the provincial governors and, since 2008, of the regional directors for environmental protection. The biological grounds for such regulation of harvesting were formulated as early as the 1970s [49]. The situation changed with the development of GIS techniques, and the character of the collection changed fundamentally as well. Snail specimens obtained in the field were replaced by virtual data, with every individual observed in the field being located by GPS. Shells for measurement and for supplementing the collection were gathered less frequently and in smaller numbers. At the same time the metadata base grew steadily, becoming an indispensable research tool for the biology and ecology of the Roman snail. Extensive surveys in several provinces refined the data on the distribution of local populations of this mollusc and at the same time made it possible to set limits on its commercial harvesting in each of them [50,51] (Table 3). The way in which the boundaries of local populations are delimited on the basis of the outermost occurrences of individuals is illustrated in Figures 8 and 9. This significantly raised the scientific value of the collection.

**Figure 8.**
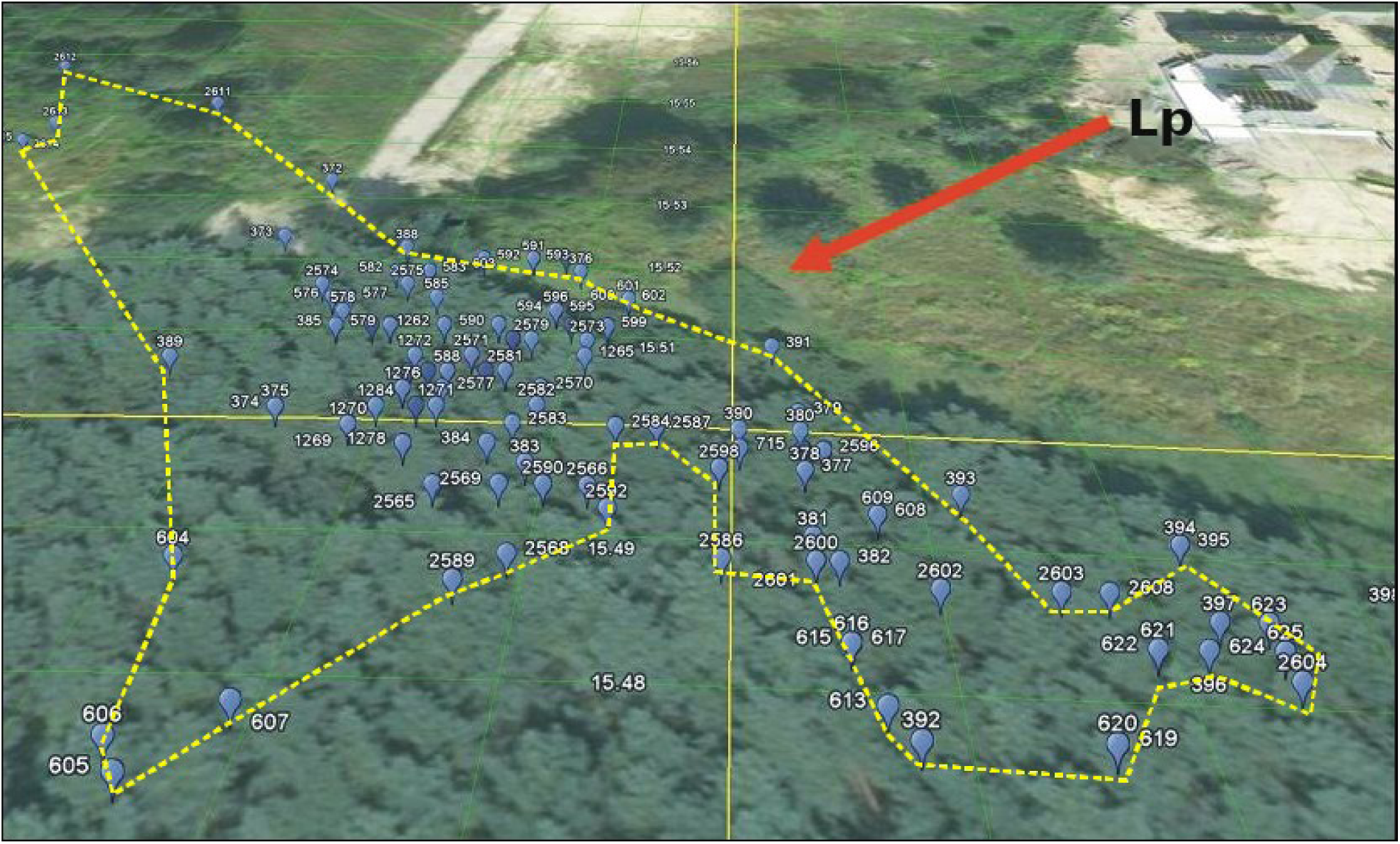
Delimitation of the boundaries of a single local population on the basis of the outermost occurrences of individuals in the population: Lp - local population. Basemap: Google Earth, imagery © Google.

**Figure 9.**
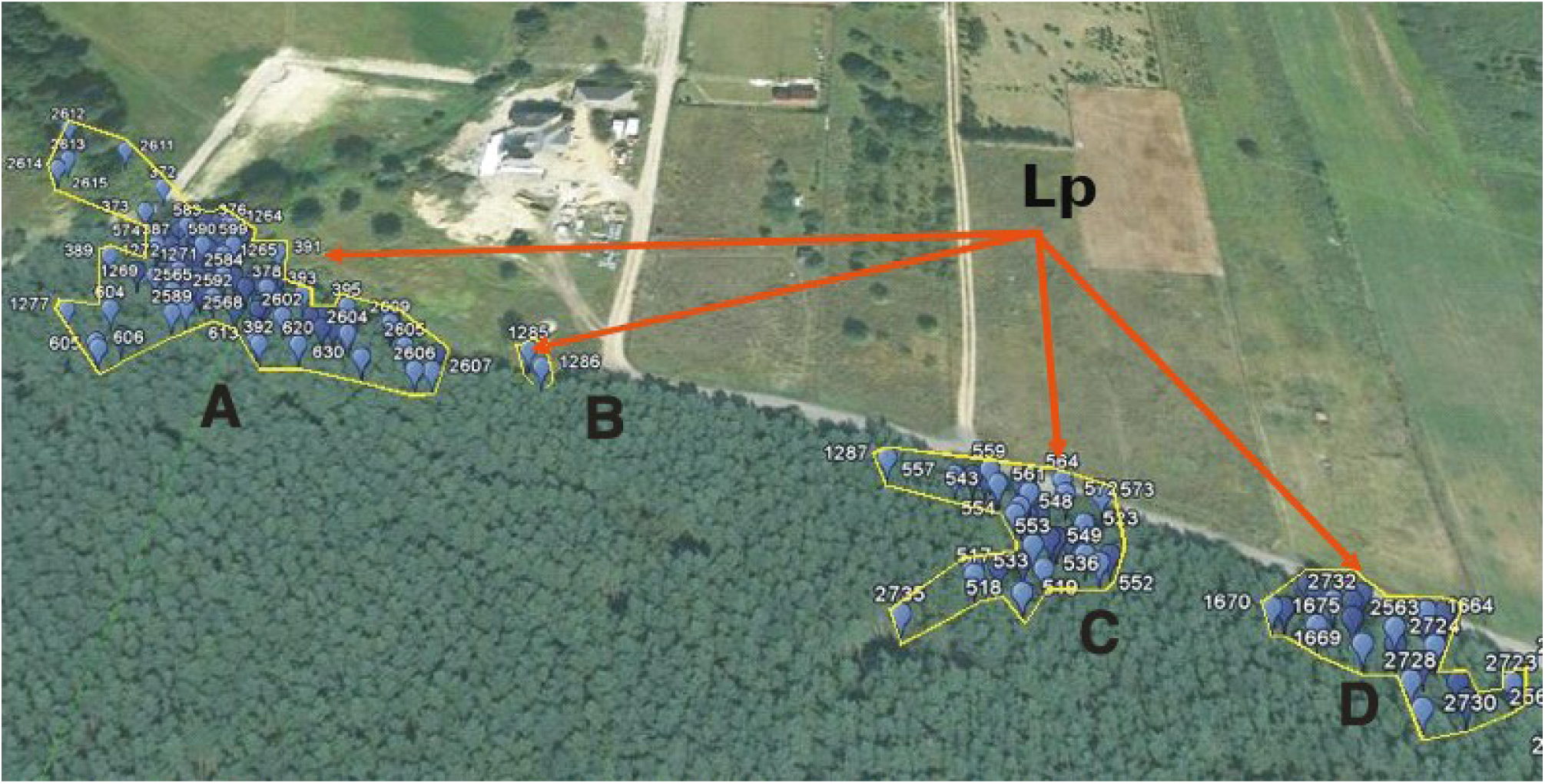
Delimitation of local populations on the basis of the outermost occurrences of individuals in the population: Lp - local populations, A-D - the local populations delimited. Basemap: Google Earth, imagery © Google.

Enriching the metadata with marked specimens observed over a longer period makes it possible to use the data set for innovative metapopulation research [52,53]. The use of GIS tools, in turn, allows the movements of a snail to be tracked precisely and the area penetrated by a single individual to be delimited (Figure 10), which was previously all but impossible. The scientific value of the collection, and especially of the metadata associated with it, is thus growing continuously.

**Figure 10.**
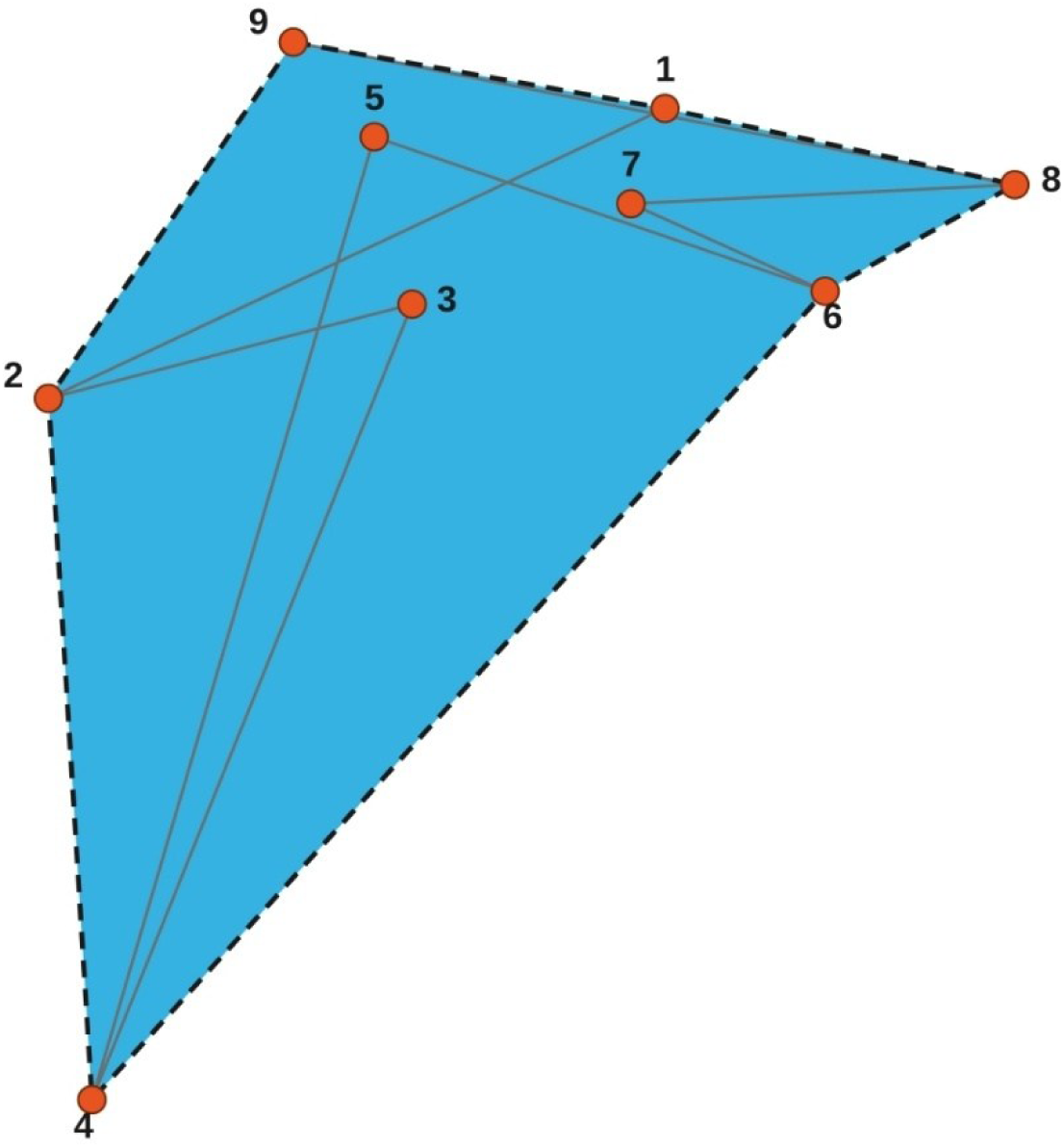
Delimitation of the boundaries and of the area penetrated by a single individual: 1-9 successive observations

#### Untapped potential

This collection, too, holds a resource that has not yet been activated. A shell is a carbonate record of the conditions under which it formed, and the composition of oxygen and carbon isotopes in the successive growth increments of land snail shells is sometimes used as an indicator of temperature and humidity. The series assembled from dated sites span more than half a century, which makes them ready material for such analyses. The preserved bodies and the shell material itself may also serve as a source of DNA for phylogeographic studies, allowing the genetic structure of populations from decades ago to be compared with the present state. Material from before the GPS era is described by verbal locality, so its retrospective georeferencing would join the two layers of the collection into a single temporal series and would make it possible to assess which of the historical sites have survived the transformation of the landscape. Publication of the records in GBIF, in turn, would make data on the distribution of a species listed in Annex V of the Habitats Directive available to the community engaged in its conservation at a European scale.

The trajectory of this collection is in a sense the reverse of the preceding one: the starting point was a set of physical specimens and the end point a resource of spatial data in which the specimen has become a supplementary rather than a primary element. Despite this reversal, both illustrate the same mechanism. Value grew not because material was being added, but because successive layers of information were being added to existing holdings: measurements, coordinates, the marking of individuals and repeated observations. In the categories of Section 4 this was, successively, enriching the data record and increasing visibility. In the categories of Section 3, what grew above all was data integrity and documented and potential research applications, supported by the ethical and legal compliance that follows from the collection being tied to the permit system of the Regional Directorates for Environmental Protection.

#### What the two examples show

Taken together, the two examples cover all five dimensions of scientific value. The collection of soil samples illustrates above all epistemic distinctiveness and unique taxonomic status; the conchological collection illustrates data integrity and the ethical and legal compliance that follows from the collection being tied to a permit system; and both document the dimension of documented and potential research applications. They also show two different routes to the same result: through the expansion of the derivatives of a set of specimens, and through shifting the emphasis from the specimen to the observation record.

## 6 Conclusions

Natural history collections today function as research infrastructure rather than as a closed archive. What decides this is the asymmetry between the broadly constant cost of maintaining the holdings and the spectrum of their applications, which widens with every new method of working on historical material. Specimens once prepared solely for the purposes of morphology now sometimes serve as a source of molecular, isotopic and phylogeographic data, provided that storage conditions and the quality of documentation allow it.

The article proposes an analytical framework that makes it possible to describe this value, to order it and to shape it deliberately. The five dimensions of scientific value (epistemic distinctiveness, data integrity, research applications, taxonomic status, and ethical and legal compliance) form a profile of a collection far more informative than any directly measured single figure such as the number of specimens. The typology of four categories of action (enriching the data record, enriching the physical specimen, establishing formal status, and increasing visibility and interoperability) converts that profile into a practical development plan that can be implemented in stages and adjusted to available resources.

The case study of the collections deposited in the Natural History Collections of AMU shows that neither the framework nor the typology is an abstraction. The same collection of soil samples that served for decades as a source of material for morphological and faunistic research has also become the basis of a collection of nomenclatural types, of a resource of SEM images, of a database of molecular sequences and of a potential node in the global infrastructure of biodiversity data. This entire expansion took place on material already deposited, without supplementing the holdings with new field acquisitions. The second collection discussed, the conchological collection of the Roman snail, followed the reverse path: from a set of shells to a resource of spatial data in which the specimen has become a supplement to the observation record. Both illustrate the same mechanism of value accruing through the addition of successive layers of information to material already held.

The framework proposed has limitations that are worth stating explicitly. It was derived from review literature and tested on two collections from a single institution, both connected with the research base of the authors, so the selection of examples is purposive rather than random. Both are zoological collections of invertebrates. The framework has not been tested on a herbarium, an osteological collection or a palaeontological collection, where the weight of the individual dimensions may be distributed differently. We also deliberately refrain from proposing a scale or a numerical indicator, which distinguishes this approach from the attempts at quantitative formalisation invoked in Section 2, including our own method for determining the relativistic value of a collection [17]. We treat the two approaches as complementary: the former concerns an aggregate assessment of the state of the holdings, the latter the identification of directions for their development. Verification of the usefulness of the framework by curators outside the authors’ circle remains an open task.

The practical consequence of the framework and the typology is that a collection development plan can be formulated in terms of concrete, accountable actions rather than general declarations about the need to protect holdings. For a curator this makes it possible to order priorities by the ratio of effort to expected gain in value; for a funding body it provides a clear criterion for assessing applications concerning collection infrastructure. In this sense the second life of a collection is not a one-off event but the result of a repeatable decision to look at the material already held with fresh eyes and with new methods.

